# Combination fedratinib and venetoclax has activity against human B-ALL with high FLT3 expression

**DOI:** 10.1101/2023.06.07.544058

**Authors:** Sean P Rinella, Haley C Bell, Nicholas J Hess, Nguyet-Minh Hoang, Thao Trang Nguyen, David P Turicek, Lei Shi, Lixin Rui, James L LaBelle, Christian M Capitini

## Abstract

Treatment of relapsed/refractory B cell acute lymphoblastic leukemia (B-ALL) remains a challenge, particularly in patients who do not respond to traditional chemotherapy or immunotherapy. The objective of this study was to assess the efficacy of fedratinib, a semi selective JAK2 inhibitor and venetoclax, a selective BCL-2 inhibitor, on human B-ALL using both single-agent and combinatorial treatments. The combination treatment of fedratinib and venetoclax improved killing of the human B-ALL cell lines RS4;11 and SUPB-15 in vitro over single-agent treatments. This combinatorial effect was not detected in the human B-ALL cell line NALM-6, which was less responsive to fedratinib due to the absence of Flt3 expression. The combination treatment induces a unique gene expression profile relative to single-agent treatment and with an enrichment in apoptotic pathways. Finally, the combination treatment was superior to single agent treatment in an in vivo xenograft model of human B-ALL, with a two-week treatment regimen significantly improving overall survival while inducing CD19 expression. Overall, our data demonstrates the efficacy of a combinatorial treatment strategy of fedratinib and venetoclax against human B-ALL expressing high levels of Flt3.

**Statement of Implication:** The combination treatment of fedratinib and venetoclax has activity against human B-ALL with high FLT3 expression and has the potential to be a salvage therapy for patients with relapsed/refractory B-ALL.

## Introduction

While significant strides have been made to treating B cell acute lymphoblastic leukemia (B-ALL) with now over 80% children cured, patients who relapse after initial treatment suffer a poor prognosis (1,2). While many relapsed/refractory pediatric patients are salvaged with CD19-directed immunotherapies like blinatumomab or tisagenlecleucel, the 12-month event free survival is only 50%, with CD19 loss representing a major barrier to the long-term success of these immunotherapies (3). It is thus essential to investigate novel combinatorial treatments with targeted pharmaceuticals to improve patient outcomes and quality of life.

The dysregulation and/or activating mutations in the protein Janus kinase 2 (JAK2) lead to downstream phosphorylation and activation of signal transducer and activator of transcription (STAT) proteins that are predominant in many cancers (4,5). In a recent Children’s Oncology Group (COG) study (6) genomic alterations involving activation of the JAK/STAT pathway were found in 22% of high-risk B cell ALL patient samples, including Philadelphia (Ph)-like B-ALL.

The BCL-2 family of proteins are central regulators of apoptosis. BCL-2 inhibits apoptosis by the preservation of mitochondrial membrane integrity, specifically preventing the pro-apoptotic proteins BAX and BAK from release into the mitochondria membrane and the subsequent release of cytochrome C (7). Furthermore, some high-risk subsets of B-ALL, such as KMT2A-rearranged B-ALL or B-ALL with hypodiploidy, express high levels of BCL-2 to prevent apoptosis (8). We and others have previously shown that dual targeting of the JAK/STAT pathway and BCL-2 can treat acute myelogenous leukemia (9) and T cell ALL (10), however the impact of targeting both these pathways in B-ALL is unknown.

The overall goal of this study was to investigate the direct anti-leukemic activity of the JAK2 inhibitor fedratinib and the BCL-2 inhibitor venetoclax either as a single-agent treatment or combinatorial treatment. By inhibiting both of these pathways at the same time, we aim to enhance their anti-leukemic effects on B-ALL.

## Materials and Methods

### B-ALL cell lines

NALM-6 (B-ALL), RS4;11 (KMT2A-rearranged B-ALL) and SUP-B15 (Ph-like B-ALL) were purchased from ATCC. NALM-6 and RS4;11 were cultured in RPMI 1640 (Corning) supplemented with 10% fetal bovine serum (FBS) (GeminiBio), 2 mmol/L L-glutamine (Gibco), and 100 mg/mL streptomycin/100 U/mL penicillin (Lonza). SUP-B15 cells were cultured in McCoy 5A (modified) media (Gibco) supplemented with 20% FBS, 100 mg/mL streptomycin and 100 U/mL penicillin. RS4;11 cells were transduced to express the luminescent reporter luciferase using pGL4.51[luc2/CMV/Neo] vector (Promega). Cell authentication was performed using short tandem repeat analysis (Idexx BioAnalytics, Westbrook, ME) and per ATCC guidelines using morphology, growth curves, and *Mycoplasma* testing within 6 months of use (InVivoGen). Cell lines were maintained in culture at 37°C in 5% CO_2_.

### Fedratinib and Venetoclax

Venetoclax (1 mL*10 mM in DMSO) and powdered compound were purchased from TargetMol. Fedratinib (1 mL*10 mM in DMSO) was purchased from Selleckchem and powdered was purchased from MedChemExpress. All concentrations for in vitro studies were made at 2x or 4x for combination studies and diluted in RPMI with 10% FBS for plating assays. Cells were treated with a range of doses ranging from 1nM to 10uM.

### Live/dead discrimination assays

Cells were collected after a 48 hour incubation with either vehicle control, fedratinib, venetoclax, or combination and live/dead discrimination was assessed through flow cytometry. Samples were labeled with Ghost Dye Red 780 (Tonbo Biosciences) and analyzed via an Attune NxT Flow Cytometer (ThermoFisher) with analysis on FlowJo V10.

### Proliferation assays

Cells were collected after a 72 hour incubation with either vehicle control, fedratinib, venetoclax, or combination using the CellTiter 96 AQueous One Solution Cell Proliferation Assay (Promega). Plates were read using a CLARIOStar microplate reader (BMG Labtech).

### RNA seq

RS4;11 cells were treated for 24 hours in triplicate (DMSO, fedratinib alone, venetoclax alone, or combination). All RNA isolation and sequencing was performed by the Gene Expression Center at the University of Wisconsin-Madison. RNA was isolated with a RNeasy kit (Qiagen) prior to a quality control check using an Agilent Bioanalyzer and NanoDrop One spectrophotometer. mRNA libraries were prepared using the TruSeq stranded mRNA kit and sequenced on an Illumina NovaSeq 6000 platform to 30 million reads per sample. Sample reads were processed by the University of Wisconsin Biotechnology Center.

### RNA-Seq analysis

Bioinformatic analysis of transcriptomic data adhere to recommended ENCODE guidelines and best practices for RNA-Seq. Alignment of adapter-trimmed (Skewer v0.1.123; (11)) 2x150 (paired-end; PE) bp strand-specific Illumina reads to the *Homo sapiens*GRCh38 genome (assembly accession NCBI:GCA_000001405.25) was achieved with the Spliced Transcripts Alignment to a Reference (STAR v2.5.3a) software (12), a splice-junction aware aligner, using annotation provided by Ensembl release 104. Expression estimation was performed with RSEM v1.3.0 (RNASeq by Expectation Maximization; (13)). To test for differential gene expression among individual group contrasts, expected read counts obtained from RSEM were used as input into edgeR (v3.16.5; (14)). Inter-sample normalization was achieved with the trimmed mean of M-values (TMM; (15)) method. Statistical significance of the negative-binomial regression test was adjusted with a Benjamini-Hochberg FDR correction at the 5% level (16). Prior to statistical analysis with edgeR, independent filtering was applied and required genes to have a count-per-million (CPM) above *k* in *n* samples, where *k* is determined by minimum read count (10 reads) and by the sample library sizes where *n* is determined by the number of biological replicates in each group. The validity of the Benjamini-Hochberg FDR multiple testing procedure was evaluated by inspection of the uncorrected p-value distribution.

Gene Set Enrichment Analysis (GSEA) software MSigDB_2023.1 was used to test for enriched pathways in the combination group compared to the other three experimental groups. A total of 1000 permutations were chosen using the Ensembl_Human_Gene_ID platform and tested against the Hallmark, KEGG and Reactome data sets, with relevant significantly enriched pathways shown (17,18). The GSEA and MSigDB are available under a Creative Commons style license and is a joint project of UC San Diego and the Broad Institute.

### Apoptosis and cell cycle assays

RS4;11 cells were treated for 24 hours (DMSO, fedratinib alone, venetoclax alone, or combination) prior to both analyses. BrdU-7AAD and Annexin V-PI analyses were performed according to manufacturer’s protocols (BD Biosciences) and analyzed via an Attune NxT Flow Cytometer (ThermoFisher, Waltham, MA) with analysis on FlowJo v10 (Ashland, OR).

### BH3 Profiling Assay

BH3 profiling is a functional assay that measures apoptotic priming of cells of interest after pre-treatment with a drug. The BH3 profiling assay measures the extent of mitochondria outer membrane permeability that occurs in response to pro-apoptotic BH3 peptides, which mimic the activity of pro-apoptotic BH3 only proteins from the BCL-2 family methods. Methods have been previously described by others(19).

### In vivo studies

NOD/SCID/*Il2rg^tmwjl^*/Szj (NSG) breeder mice were purchased from Jackson Labs (Bar Harbor, ME) and bred internally at the Biomedical Research Model Services (BRMS) Breeding Core at the University of Wisconsin-Madison in accordance with the Guide for the Care and Use of Laboratory Mice and experiments were performed under an IACUC approved animal protocol (M005915). Male and female NSG mice were used at 8-16 weeks of age and injected via tail vein with 2 x10^6^ RS4;11 cells on Day 0. Peripheral blood was obtained via retro-orbital bleeds conducted on Days 10 (engraftment), Day 21 (pre-treatment), and Day 35 (post-treatment). Mice were treated with fourteen days of once daily treatment with vehicle, fedratinib, venetoclax, or combination starting at Day 21 (5 days on, 2 days off). Venetoclax was prepared for in vivo use by dissolving drug into the following: 5% DMSO, 30% PEG400, 7.5% Tween20, 7.5% Tween80 and 50% sterile water. Fedratinib was prepared for in vitro use by dissolving drug into 100% sterile water. Venetoclax was administered via intraperitoneal (IP) injections at a dose of 6.25 mg/kg. Fedratinib was administered via oral gavage at a dose of 100 mg/kg. Mice were monitored for survival and percent weight change and euthanized when a humane endpoint was reached. Peripheral blood was treated with ACK lysis buffer (Lonza) and then stained in flow buffer (PBS, 10% FBS) with CD19 (HIB19), CD20 (2H7), CD22 (HIB22), PD-L1 (MIH2), FAS (DX2), HLA-DR (L243), BCL2 (100) and BAX (2D2), all purchased from Biolegend, prior to quantification on an Attune NxT Flow Cytometer with analysis on FlowJo V10.

### Statistical analysis

Statistics were performed using GraphPad Prism version 9.0 for the Macintosh OS (GraphPad Software). Data were expressed as mean ± SEM. For analysis of three or more groups, a non-parametric one-way ANOVA test was performed with the Bonferroni or Sidak’s multiple comparisons post-test. Analysis of differences between two normally distributed test groups was performed using a two-sided Mann Whitney test. A p-value less than 0.05 was considered statistically significant.

### Data availability statement

By the time of publication all RNA seq data will be reposited in the Gene Expression Omnibus (GEO). All other data will be made available upon reasonable request.

## Results

### Combinatorial treatment of fedratinib and venetoclax effectively kills human B-ALL in vitro

To assess the efficacy of JAK inhibitors to kill human B-ALL cells, we tested ruxolitinib (JAK1/2 inhibitor) and fedratinib (semi-selective JAK2 inhibitor) on the human B-ALL cell line RS4;11. Cells were treated with a range of doses from 1nM to 10uM for 48 hours prior to analysis for cell survival (Figure 1A-B). Cells were also treated with single agent venetoclax (Figure 1C). Based on the dose-curves, the calculated IC50 for fedratinib was 600nM but the IC50 for ruxolitinib was not reached.

**Figure 1.**
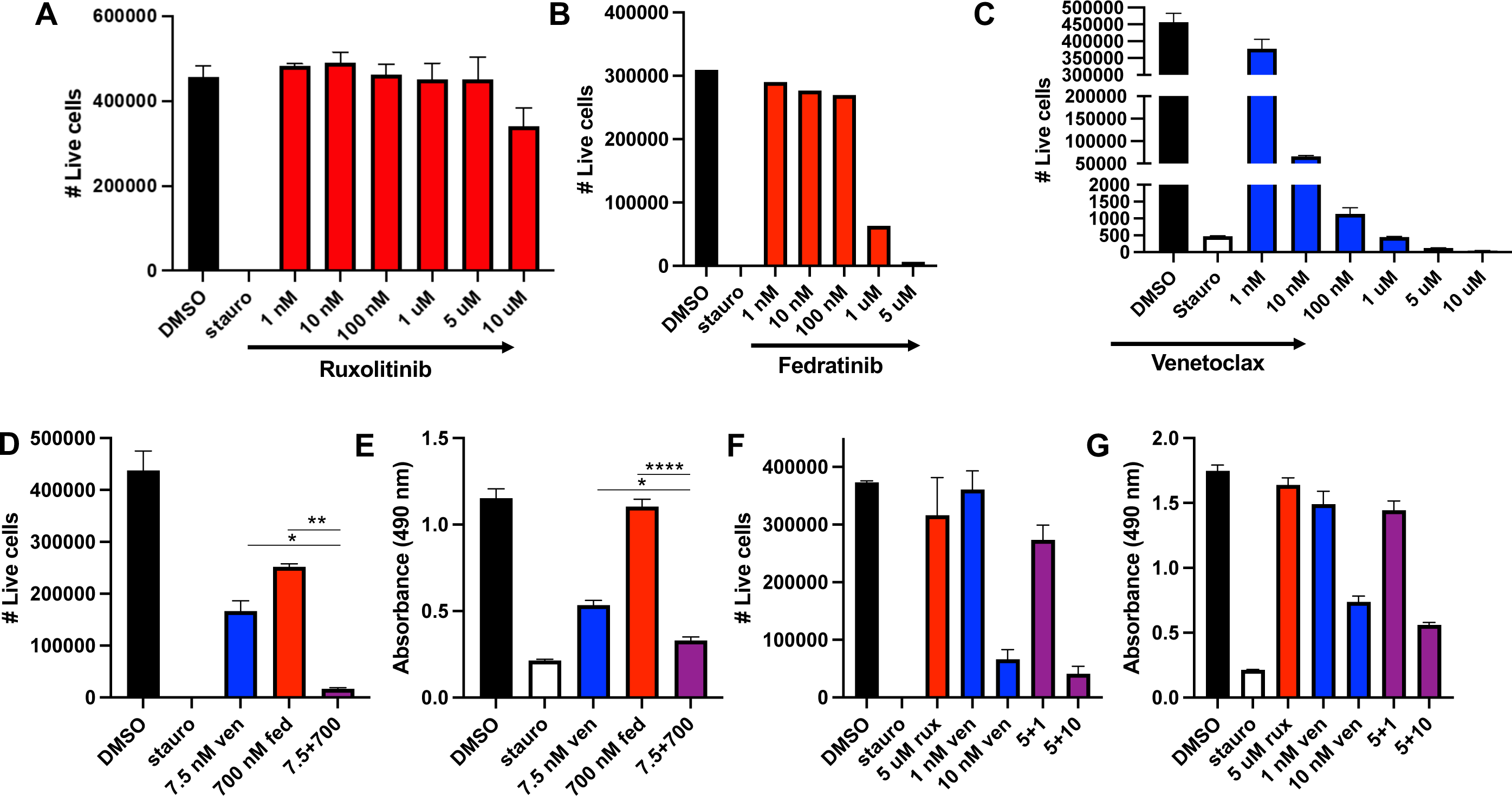
Fedratinib, but not ruxolitinib, combines with venetoclax to improve killing of human B-ALL. The human B-ALL cell line RS4;11 was treated with a dose curve of single agent ruxolitinib (A), fedratinib (B) or venetoclax (C) and number of live cells quantified by flow cytometry after 48hrs. (D-G) RS4;11 cells were treated with the indicated concentration of fedratinib and venetoclax (D-E) or ruxolitinib and venetoclax (F-G) for 48 or 72hrs. Live cells were quantified by flow cytometry and proliferation rate measured by colorimetric MTS assay. Fed = fedratinib. Rux = ruxolitinib. Ven = venetoclax. * = p < 0.05, ** = p < 0.01, **** = p < 0.0001

We next wanted to explore whether or not fedratinib would be more efficacious when combined with venetoclax. To test this, we treated RS4;11 with single agent fedratinib at 700nm, single agent venetoclax at 7.5nM and the combinatorial treatment (Figure 1D-E). Combinatorial treatment was significantly more effective than either single agent treatment and similar to the positive control staurosporine. We next tested if ruxolitinib had a similar beneficial effect as fedratinib (Figure 1F-G). Ruxolitinib did not improve the killing of RS4;11 over single-agent venetoclax (Figure 1F-G), even at higher doses. To ensure that the combinatorial effect of fedratinib and venetoclax was not specific to RS4;11, we tested both the single agent and combinatorial treatments on the human B-ALL cell lines SUPB-15 and NALM-6. The combinatorial treatment was equally effective on SUPB-15 (Figure 2A-B) but surprisingly, the combinatorial treatment did not improve the killing of NALM-6 cells compared to venetoclax alone (Figure 2C-D). These data suggest that while the combinatorial treatment of fedratinib and venetoclax can be highly effective in human B-ALL, there are a subset of B-ALLs that remain resistant to this combinatorial treatment.

**Figure 2.**
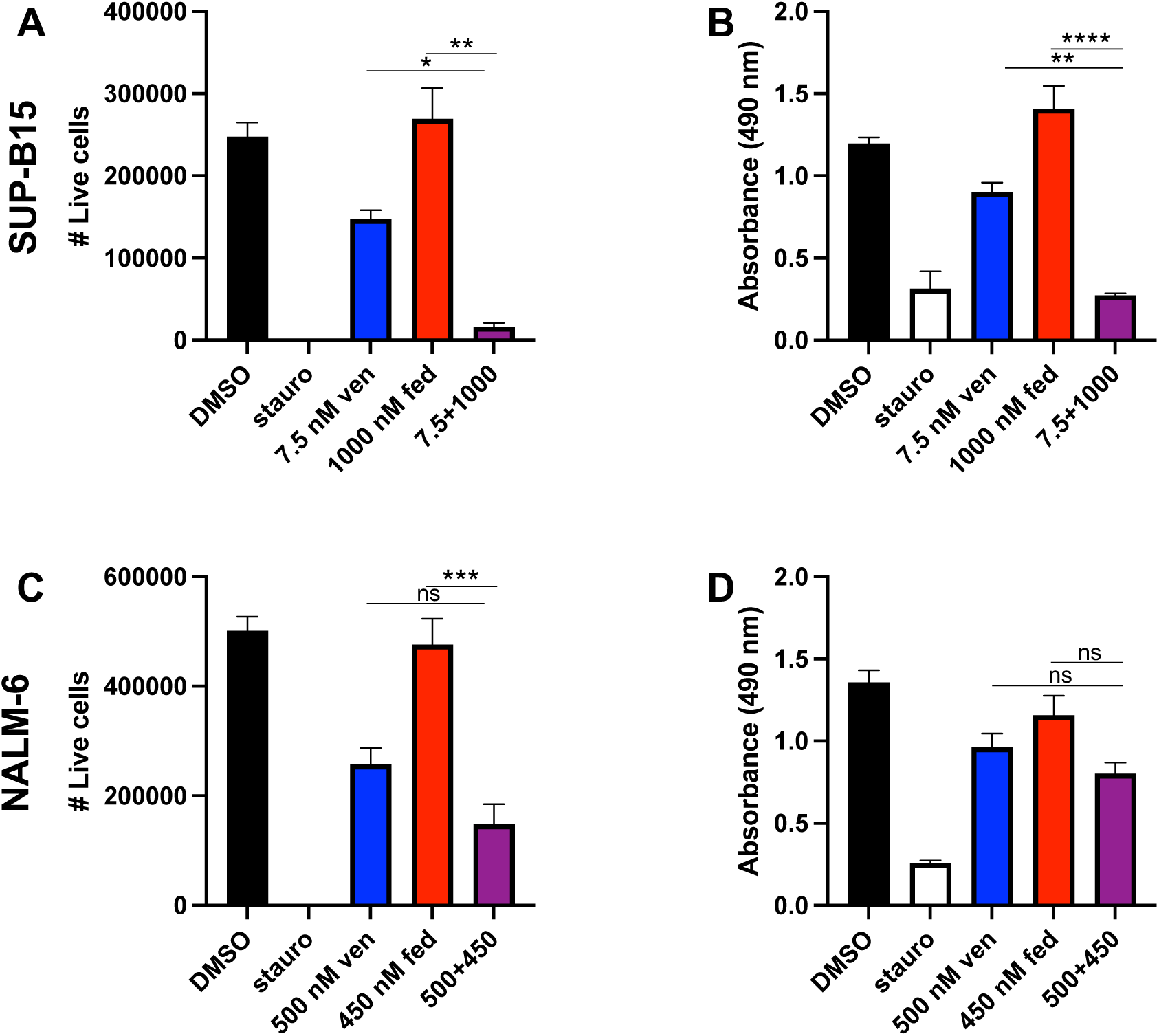
Combinatorial treatment of venetoclax and fedratinib is efficacious in the human B-ALL cell line SUP-B15, but not NALM-6. Human B-ALL cell lines NALM-6 and SUP-B15 were treated with venetoclax in combination with fedratinib. Live cells were quantified by flow cytometry as previously described (A,C) and proliferation was assessed by an MTS assay as previously described (B,D). ns = not significant, * = p < 0.05, ** = p < 0.01, *** = p < 0.001, **** = p < 0.0001

### Resistance to fedratinib treatment is correlated with absence of Flt3 expression

To determine the precise activity of and contribution to cell killing by fedratinib and venetoclax, we performed several additional dose-escalation studies on all three human B-ALL cell lines. Keeping a constant venetoclax dose of 5 nM, both RS4;11 and SUPB-15 experienced significant cell death that was further enhanced with increasing doses of fedratinib. Conversely, NALM-6 was mostly refractive to venetoclax at 5 nM with increasing concentrations of fedratinib modestly improving the killing efficiency (Figure 3A-C, left panels). Using a constant dose of fedratinib of 525 nM, increasing doses of venetoclax was able to significantly increase cell death in both RS4;11 and SUP-B15 while NALM-6 remains refractory as before (Figure 3 A-C, right panels).

**Figure 3.**
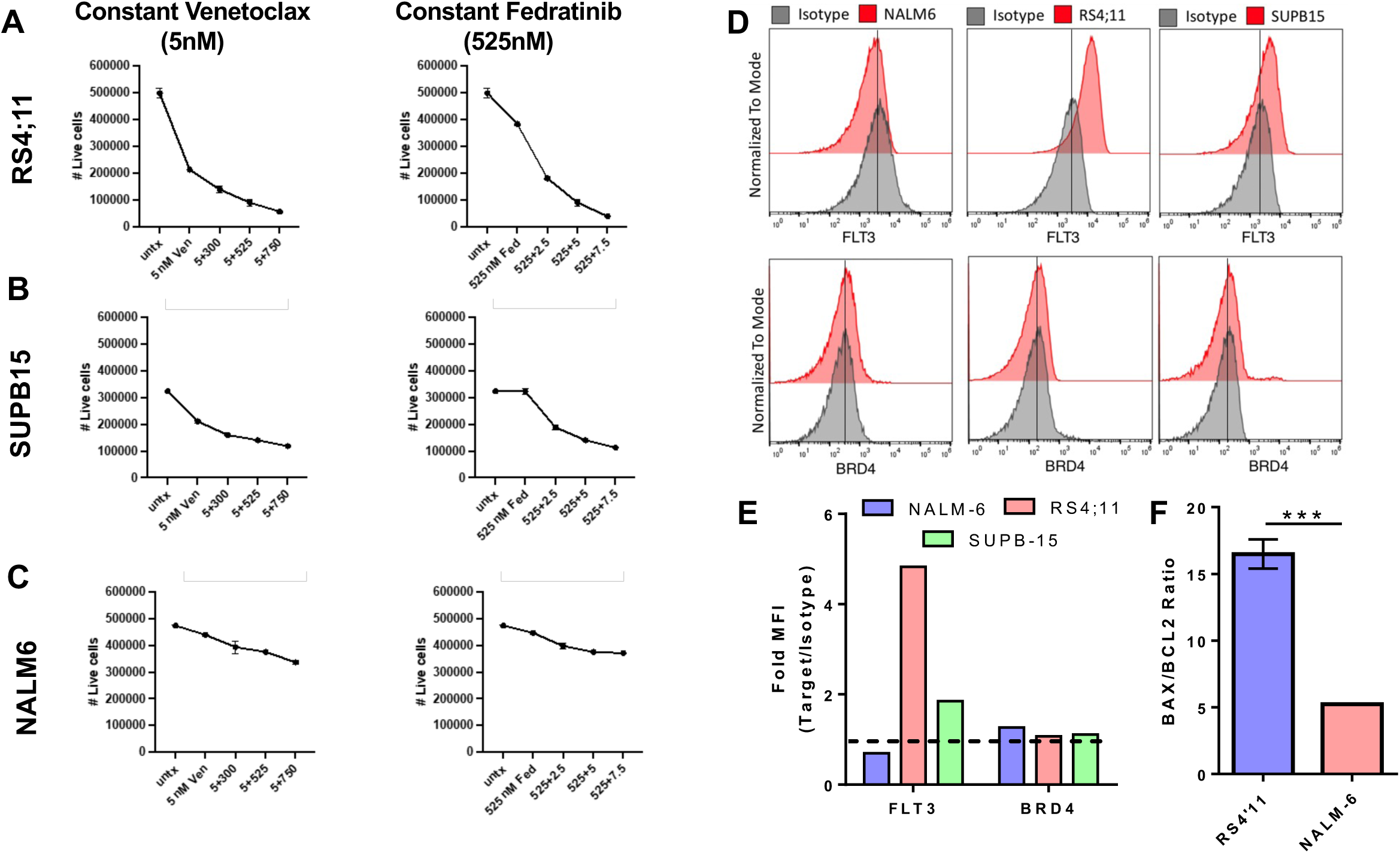
Fedratinib activity is associated with Flt3 expression. Combination dose curves were performed on RS4;11 (A), SUP-B15 (B) and NALM-6 (C), keeping either the venetoclax dose (left panel) or fedratinib dose (right panel) constant to determine the potency of the combinatorial treatment on different human B-ALL cell lines. (D-F) Expression of both Flt3 and Brd4 (D-E) and the ratio of BAX to BCL2 (F) were measured in each cell line by flow cytometry. Fold MFI represents the median fluorescent intensity of the marker divided by the isotype control. *** p < 0.001.

To assess a potential mechanism for NALM-6 resistance to both venetoclax and fedratinib treatment, we measured the expression of downstream targets BCL-2, BRD4 and FLT3 in each human B-ALL cell line. Notably, NALM-6 had no expression of FLT3, which is also a target of fedratinib while RS4;11, which has the greatest sensitivity to fedratinib, had the highest expression of Flt3 (Figure 3D-E). We also observed that NALM-6 was more resistant to venetoclax treatment than either RS4;11 or SUPB-15 (Figure 1, 2). Interestingly, NALM-6 had a significantly lower ratio of the pro-apoptotic protein BAX to the anti-apoptotic protein BCL2 relative to RS4;11, which correlated with venetoclax sensitivity (Figure 3F).

### Combination therapy differentially enriches pathways related to cell death

Because there are many potential downstream targets impacted by both fedratinib and venetoclax, we wanted to identify the dominant pathways altered in RS4;11 post single-agent and combination treatment via bulk RNA sequencing. RS4;11 cells were plated similarly as described above and exposed for 24 hours to fedratinib (525 nM), venetoclax (5 nM), or combination and compared to untreated cells. Cells were harvested after 24 hours for RNAseq. Whole transcriptome analysis revealed that each treatment regimen induced a unique gene expression profile (Figure 4A-B). Comparing each treatment regimen to untreated cells, fedratinib had the fewest number of significantly different genes while the combinatorial treatment had the most (Figure 4C-E). Interestingly in all cases, most of the differentially expressed genes were down-regulated and not up-regulated as a result of treatment (Figure 4C-E). We also completed a gene set enrichment analysis (GSEA) that identified pathways associated with apoptosis, DNA repair and proliferation predominantly altered in the combinatorial treatment versus all three other conditions (Figure 4F). A heatmap of the differentially enriched genes primarily impacting the enrichment into the apoptotic pathway is also shown (Figure 4G). Functional assays of both apoptosis and cell cycle were consistent with RNA-seq analyses. A higher fraction of cell death was observed in combination treatment in RS4;11 cells compared to single agent fedratinib or venetoclax (Figure 5A). There also was evidence of G2-M arrest in combination treatment compared to single agent treatment (Figure 5B). Finally, BH3 profiling suggests venetoclax differentially contributes to BIM-mediated apoptosis (Figure 5C-G).

**Figure 4.**
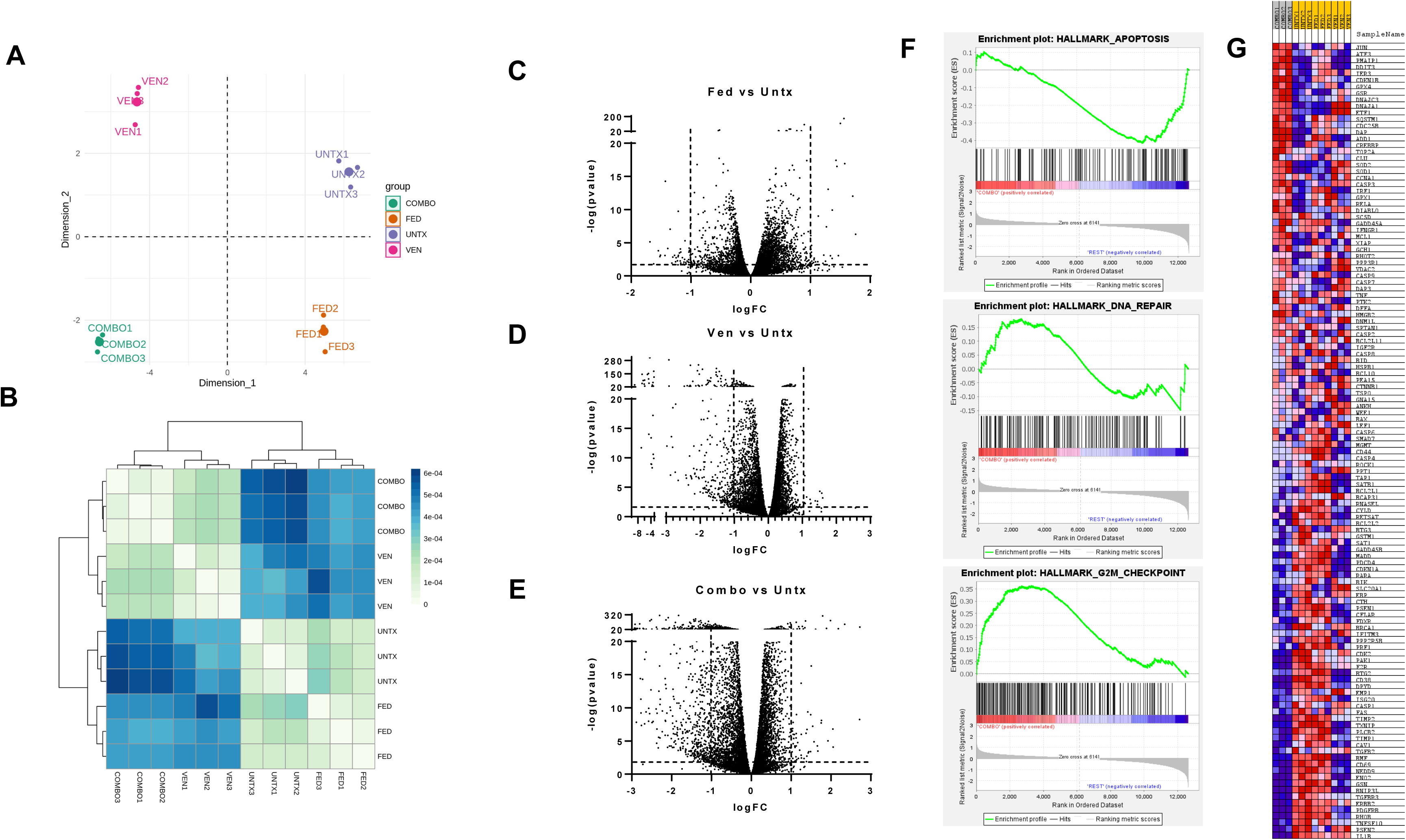
Combination venetoclax and fedratinib treatment induces a unique gene expression profile. RS4;11 cells were left untreated or treated with 5 nM ventoclax, 525 nM fedratinib or the combination treatment for 24 hrs prior to RNA isolation and bulk sequencing. (A-B) Transcriptome analysis of each treatment group by two-dimensional multidimensional scaling (MDS) (A) and a clustered image map reveals each treatment group induces a unique transcriptomic profile. (C-E) Volcano plots showing the number of positively and negatively regulated genes in each treatment group relative to the untreated control. (F) Gene set enrichment analysis of the combination treatment against all other treatments was performed with three gene sets shown. (G) Heatmap showing the genes contributing to the enrichment score of the apoptosis gene set.

**Figure 5.**
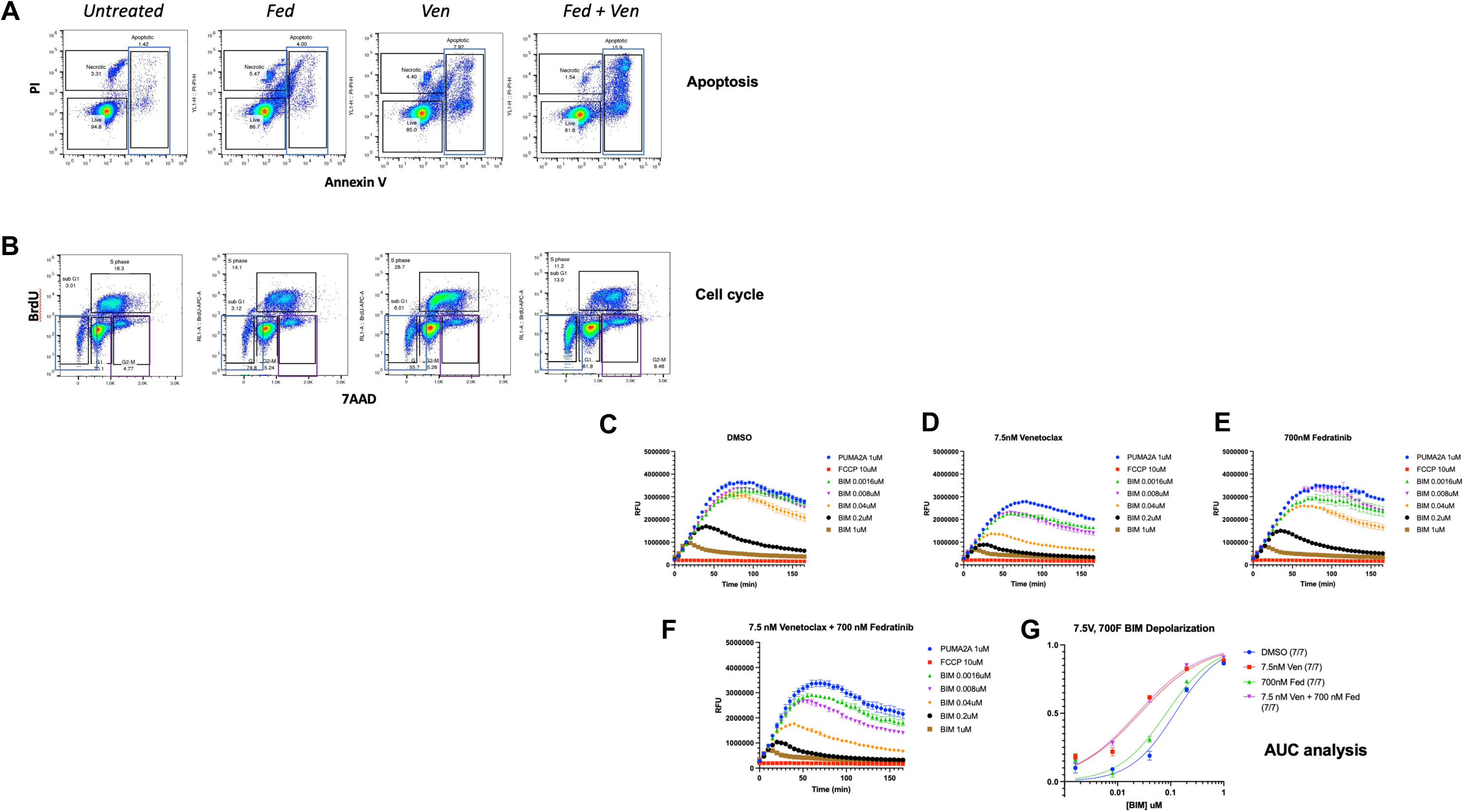
Apoptosis, cell cycle and BH3 profiling is dysregulated after combinatorial treatment with venetoclax and fedratinib. Flow cytometric assay of apoptosis (A) and cell cycle (B) after treatment with (need doses and target cell line). BH3 profiling assay examining BIM depolarization after exposure to (C) DMSO control, (D) 7.5 nM venetoclax, (E) 700 nM fedratinib and (F) combination. (G) AUC analysis of depolarization BH3 profiling curves in RS4;11 B-ALL.

### Combination therapy reduces leukemic burden and improves overall survival in RS4;11 derived xenograft model of B-ALL

Based on the efficacy of fedratinib and venetoclax observed *in vitro*, a xenograft model of B-ALL was established utilizing RS4;11 cells and NSG mice to test these drugs as a combination therapy *in vivo*. Our group has previously published data in T-ALL using similar combination therapy, however, unlike in T-ALL we did not see evidence of leukemia related CNS disease with the RS4:11 B-ALL model (Suppl Figure 1). NSG mice were injected with luciferase^+^ RS4;11 cells on Day 0 and then randomized to one of the following groups: DMSO, single agent venetoclax, single agent fedratinib, or combination. Peripheral blood was obtained via retroorbital bleeds on days 10 (engraftment), 21 (pretreatment), and 35 (post treatment) to assess B-ALL disease burden (Figure 6A). Treatment started on day 21 and lasted for 14 days (5 days on, 2 days off) with 6.25 mg/kg venetoclax IP and/or 100 mg/kg fedratinib via gavage.

**Figure 6.**
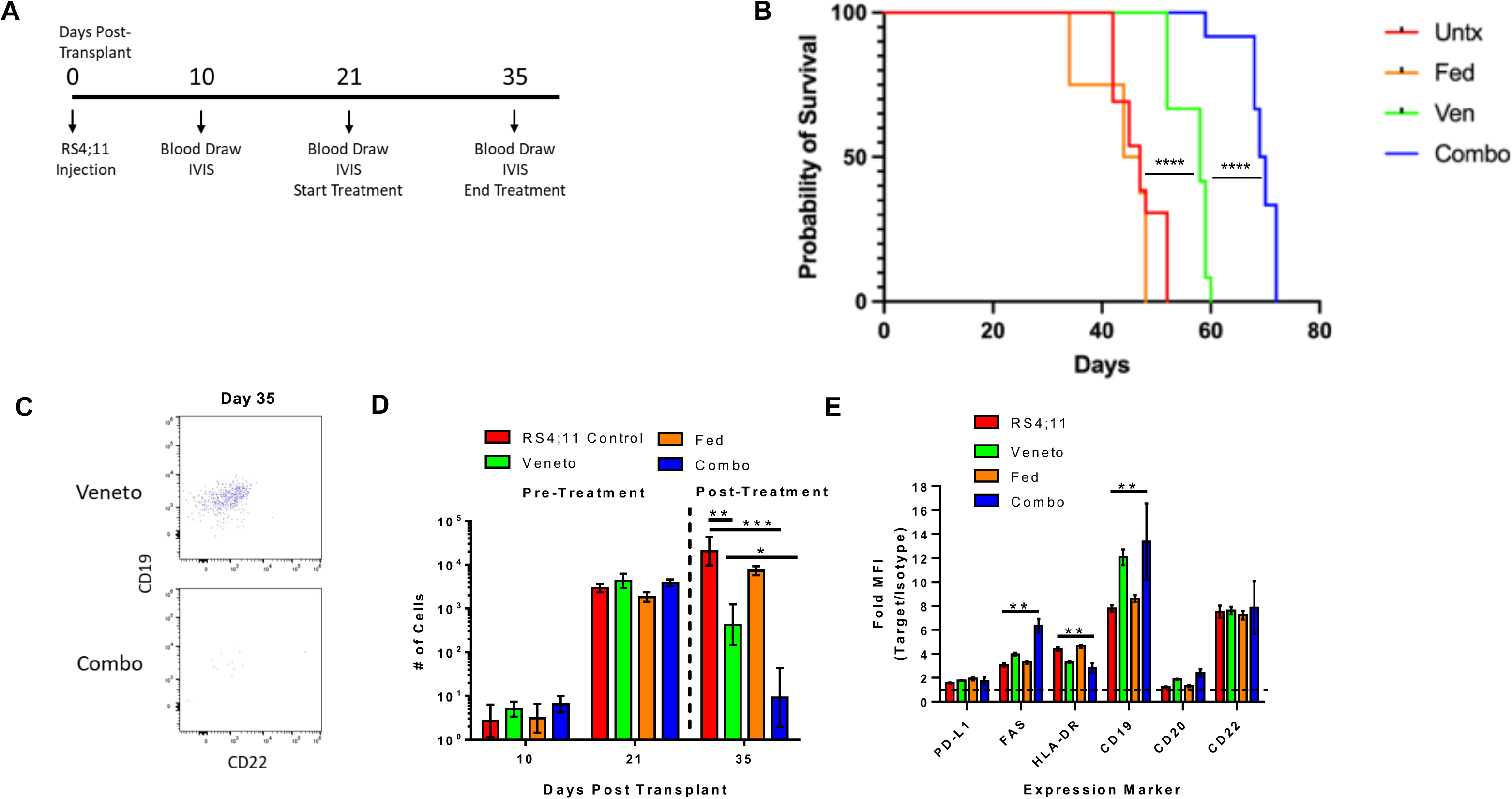
Fedratinib and venetoclax combination treatment is superior to single-agent treatment in vivo. (A) RS4;11 was transplanted into non-irradiated NSG mice via tail vein injection at 2E6 cells per mouse and allowed to engraft for 21 days prior to starting treatment. On days 21-35, Venetoclax was given IP at a dose of 6.25 mg/kg while fedratinib was given gavage at a dose of 100 mg/kg. Retro-orbital blood draws were taken at day 10, 21 and 35 to assess cancer burden. (B) Kaplan-Meier survival curve of each treatment. (C-D) Representative dot plot of RS4;11 taken from the blood at day 35 (C) and quantification across all times (D). Representative IVIS images (E) and total flux quantification of all mice at the indicated time points (F). (G) A select number of B-ALL cancer markers expression profiles were assessed at day 35 across all treatment conditions. Fold MFI represents the median fluorescent intensity of the marker divided by the isotype control. Data is representative of two independent experiments. * p < 0.05, ** p < 0.01, *** p < 0.001.

Combination treatment prolonged the survival of the mice and decreased the RS4;11 burden the most compared to the other treatment groups (Figure 6B). Venetoclax alone also significantly improved overall survival when compared to fedratinib alone and untreated mice (Figure 6B). Peripheral blood counts and IVIS imaging throughout the experiment revealed that combination treatment was highly effective at decreasing the RS4;11 burden in vivo (Figure 6C-D). Flow cytometric analysis of RS4;11 taken from the peripheral blood also showed a significant increase in MFI of FAS and CD19 in the combination group (Figure 6E). Overall, these data support the role of fedratinib and venetoclax as a combination therapy against FLT3^+^ B-ALL, with potential modulation of cell surface marker expression of CD19.

## Discussion

Treatment of relapsed/refractory B-ALL remains a challenge, particularly in patients who do not respond to traditional chemotherapy or immunotherapy. Targeting BCL-2 using venetoclax may be a viable option in subsets of B-ALL like KMT2A-rearranged and hypodiploid B-ALL. In previous work, a single dose of 12.5 mg/kg venetoclax in a RS4;11-derived xenograft model led to a maximal tumor growth inhibition of 47% and tumor growth delay of 26%. However, maximal effects were seen when combining venetoclax with chemotherapy (20). Other studies demonstrating efficacy of venetoclax in B-ALL include combining with MCL-1 inhibitors (21, 22), monoclonal antibodies such as daratumumab in relapsed/refractory B-ALL (23), AKT inhibitors (24, 25), MDM2 inhibitors (26), and also with BCR-ABL inhibitors (27). No prior studies have looked at the combination of JAK2 inhibitors with venetoclax in B-ALL. The dysregulation of JAK/STAT related genes has been previously shown to be found in high-risk patient samples of B-ALL (6). One group identified rearrangements of JAK2 in Ph like B-ALL involving the activation of mutations in IL7R and FLT3, and deletion of SH2B3, which encodes the JAK2 negative regulator LNK (28).

The goal of this study was to suppress leukemia proliferation related to JAK/STAT dysregulation while at the same time inhibiting BCL-2 to prevent escape from apoptosis in models of human B-ALL. To do this, we tested fedratinib, a semi selective JAK2 inhibitor, and venetoclax, a potent BCL2 inhibitor, to assess the efficacy of this combination in 3 models of human B-ALL in vitro and with the RS4;11 cell line in vivo. Fedratinib is FDA approved for patients with myeloproliferative disorders, primarily myelofibrosis (29, 30) and has been used to salvage patients that have failed ruxolitinib (31). Furthermore, fedratinib has been used to decrease tumor burden in Hodgkin lymphoma and mediastinal large B-cell lymphoma in vitro and in vivo (32). However, fedratinib has not previously been used for B-ALL. Analysis of BRD4 and FLT3 was included for all 3 cell lines in our study as fedratinib has been shown to target both BRD4 (33, 34) and FLT3 (33) in addition to its activity against JAK2.

Our data both in vitro and in vivo demonstrated that the combination of fedratinib and venetoclax more effectively treated B-ALL than single agent treatment. We also examined the use of another JAK2 inhibitor, ruxolitinib, which we had previously shown synergized with venetoclax to treat T cell ALL (10). Similar to observations by others (21), we found that ruxolitinib alone was ineffective in doses up to 10 uM. In contrast, combination fedratinib and venetoclax did demonstrate efficacy in the Ph+ B-ALL cell line SUP-B15, suggesting that inhibiting the JAK/STAT and BCL-2 pathways simultaneously is particularly effective in Ph-like and FLT3+ high risk subsets compared to more standard B-ALL models of disease such as NALM-6. Fedratinib also inhibits FLT3, which is commonly overexpressed in KMT2A-rearranged B-ALL (35) and was found to be highly expressed in RS4;11. The association of FLT3 and BCL2 in B-ALL is unclear but GSEA suggests changes primarily in proliferation, apoptosis and DNA repair pathways with combination fedratinib and venetoclax treatment. Functional assays of both apoptosis and cell cycle were consistent with these RNA-Seq analyses. A higher fraction of cell death was observed in combination treatment compared to single agent fedratinib or venetoclax. Furthermore, there also was evidence of G2-M arrest in combination treatment compared to single agent treatment. BH3 profiling suggests venetoclax differentially contributed to BIM-mediated apoptosis. The mitochondrial outer membrane priming of the combination therapy appears to reflect that of venetoclax alone. The addition of fedratinib, while not increasing mitochondrial sensitivity to the BIM BH3 peptide, does synergize with venetoclax to increase cell death (Figures 1, 5A, 6B) but appears to do so via a different mechanism. Further testing utilizing a FLT3 and/or BIM knock in or knock out lines could help determine the role of these molecules in response to the combination therapy.

Finally, we demonstrated efficacy of combination fedratinib and venetoclax therapy in a B-ALL xenograft model. In fact, with only a 2-week treatment we were able to identify reduced leukemic burden and an increase in overall survival benefit with combination therapy. Our group attempted to reduce potential toxicity in our in vivo model by reducing the DMSO concentration in our venetoclax drug preparation for in vivo use and gavaging with sucrose-soaked gavage tips to reduce stress on mice as much as possible, using techniques described by others (36). Our group also identified an increase in CD19 expression in peripheral blood samples obtained from mice that received either venetoclax or combination therapy. One group recently demonstrated pre-sensitizing B cell malignancies, including B-ALL, through venetoclax exposure increases cytotoxic efficacy of CD19 directed CAR-T therapy (37). These findings along with our data with combination venetoclax plus fedratinib suggest these targeted therapies may serve as a potential bridge therapy to sensitize B-ALL to CD19 targeted treatments such as blinatumomab or CAR T cells.

The development of novel treatment strategies for relapsed/refractory B-ALL is essential for the subset of patients with poor outcomes due to failure to respond to initial conventional chemotherapy. The combination of fedratinib and venetoclax is an effective treatment strategy to reduce the burden of disease, particularly in FLT3^+^ B-ALL, through reduction in proliferation and DNA repair pathways and induction of apoptosis. Combination fedratinib and venetoclax may also potentially serve as a bridge to CD19-directed therapies such as CAR T cell and future studies should explore if these therapies can help B-ALL avoid potential CD19 antigen escape.

## Acknowledgements

We would like to thank the University of Wisconsin Carbone Cancer Center (UWCCC) Flow Cytometry and Small Animal Imaging and Radiotherapy core facilities, who are supported in part through NCI/NIH P30 CA014520 as well as the UW Biotechnology Center.

## Author contributions

SPR, NJH and CMC designed the experiments, analyzed and interpreted results and wrote the manuscript; SPR, NJH, HCB, LS, NMH, LR, DPT, TN and NJH conducted experiments and analyzed data; SPR, NJH, DPT, TN and HCB generated figures; LR, JLL and CMC supervised experiments. All authors read and approved the manuscript.

## Figure Legends

**Supplementary Figure 1. FISH of bone marrow and spinal cord samples from untreated Day 51 RS4;11 transplanted NSG mice show very limited CNS infiltration during late stage leukemic disease.** Bone marrow and spinal cord tissue were harvested on untreated Day 51 transplanted RS4;11 NSG mice. Human and mouse CEP probes were used to quantify human RS4;11 cells versus mouse cells via FISH to determine disease burden in bone marrow and spinal cord tissues.

