## Supplemental Figure 1 for "Combination fedratinib and venetoclax has activity against human B-ALL with high FLT3 expression"

RS4;11

### Day 51 bone marrow

| Probe | # of cells with human CEP probe (Green) | # of cells with mouse CEP probe (Red) |
|---|---|---|
| ***Human All CEPs (G) / Mouse All CEPs (R)*** | ***196 / 200 (98.0%)*** | ***4 / 200 (2.0%)*** |

***Probe:*** ***Human CEP (G) / Mouse CEP (R)***

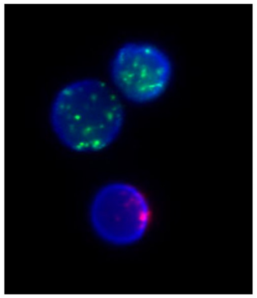

### Day 51 spinal cord

| Probe | # of cells with human CEP probe (Green) | # of cells with mouse CEP probe (Red) |
|---|---|---|
| ***Human All CEPs (G) / Mouse All CEPs (R)*** | ***10 / 200 (5.0%)*** | ***190 / 200 (95.0%)*** |

***Probe:*** ***Human CEP (G) / Mouse CEP (R)***

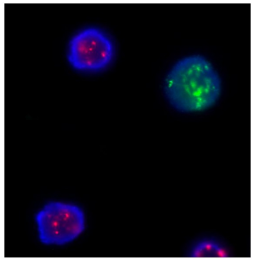

Supplemental Figure 1
